# Obligate Endosymbiosis Explains Genome Expansion During Eukaryogenesis

**DOI:** 10.1101/2022.11.17.516875

**Authors:** Samuel H. A. von der Dunk, Paulien Hogeweg, Berend Snel

## Abstract

The endosymbiosis of an alpha-proteobacterium that gave rise to mitochondria was one of the key events in eukaryogenesis. Common patterns in eukaryogenesis and present-day endosymbiotic relations include genomic streamlining of the symbiont, endosymbiotic gene transfer, and regulatory control by the host through protein targeting to the symbiont. One unique outcome for eukaryogenesis was a much more complex cell with a large genome, which may or may not be related to the mitochondrial endosymbiosis. Despite the existence of many plausible hypotheses for the observed patterns, a constructive evolutionary model in which these hypotheses can be studied is still lacking.

Here we construct an evolutionary model of cell-cycle regulation to study how obligate endosymbiosis between two prokaryote-like cells impacts cellular behavior and genome evolution. The model does not predefine an explicit fitness criterion and thereby allows for the evolution of various emergent behaviors. For instance, even though we do not allow for communication between host and symbiont, they achieve implicit cell-cycle coordination through their interaction with the environment. This evolved cell-cycle coordination can drive genome expansion as well as symmetry breaking in genome size. Many replicate runs of our evolution experiment yield organisms with a large host and small symbiont genome, but interestingly, some yield the opposite. Still on long timescales, organisms with a large host and small symbiont genome perform best, and mimic the outcome of eukaryogenesis.

By designing and studying a constructive evolutionary model of obligate endosymbiosis, we uncovered some of the forces that may drive the patterns observed in nature. Our results provide a theoretical foundation for patterns related to the mitochondrial endosymbiosis, such as genome size asymmetry, and reveal evolutionary outcomes that have not been considered so far, such as cell-cycle coordination without direct communication.

## Introduction

The mitochondrial endosymbiosis was one of the pivotal steps in the evolution of eukaryotes from prokaryotes. Yet, the sequence of events during eukaryogenesis let alone their driving forces remains largely unknown due to the lack of intermediate species between prokaryotes and eukaryotes, and the lack of evolutionary trajectories analogous to eukaryogenesis. Currently, conflicting theories have been proposed about the role of the mitochondrial endosymbiosis in the complexification of the host genome and cell. The mito-late hypothesis posits that complexity was required for endosymbiosis, such as phagocytotic machinery to engulf the endosymbiont (Kurland et al., 2006), a nucleus to protect the host genome against foreign molecules (Martijn and Ettema, 2013; Martin et al., 2015), and complex regulation and intracellular trafficking systems to control and communicate with endosymbionts (Stairs and Ettema, 2020). Conversely, the mito-early hypothesis states that mitochondria were required for complexity, to provide the energy for a large genome and cell (Lane and Martin, 2010; Schavemaker and Muñoz-Gómez, 2022).

Computational models are especially powerful for probing big questions about major evolutionary transitions where data are limited. Most hypotheses regarding the role of the mitochondrial endosymbiosis in eukaryogenesis are focused on the causes of endosymbiosis, in particular the metabolic context, but many controversies remain (e.g. see Box 1 in López-García and Moreira, 2020). However, the metabolic repertoire contributed relatively little to eukaryotic complexity (Vosseberg et al., 2021)—as opposed to the regulatory repertoire for instance—so a pure metabolic viewpoint is insufficient to understand the relation between endosymbiosis and complexity. Here we want to explore a complementary viewpoint, looking at the consequences rather than the causes of endosymbiosis by studying how obligate endosymbiosis between two prokaryote-like entities shapes the evolution of their genomes and gene regulatory networks. By designing and studying a constructive evolutionary model of cell-cycle regulation, we address the question of how endosymbiosis and complexity are related. Apart from this general question, we study with our model more specific hypotheses about the evolutionary mechanisms underlying eukaryotic complexity.

One hallmark of biological complexity that theories on eukaryogenesis seek to explain is the large eukaryotic genome. At least part of this eukaryotic genome originates from endosymbiotic gene transfer and is therefore a consequence of endosymbiosis. Several driving forces have been proposed to explain rampant endosymbiotic gene transfer from the mitochondrion to the nucleus, but the validity and quantitative contribution of each of these forces remains unresolved. For instance, endosymbiotic gene transfer could grant the host tighter regulatory control over the symbiont (Nowack, 2014), or rescue symbiont genes from Muller’s ratchet caused by high mutation rate, population bottlenecks and lack of recombination (Allen and Raven, 1996; Martin and Herrmann, 1998). Alternatively, numeric dominance of the endosymbiotic genome might be sufficient to explain how most genes end up in the nucleus through a non-adaptive ratchet-like process, except for a few special cases (Timmis et al., 2004).

In addition to gene transfer from the mitochondrion, gene duplication was an important source of eukaryotic complexity (Vosseberg et al., 2021). What drives such host complexification is unclear, in particular because no intuition can be derived from present-day endosymbiotic relations which involve a host that is already a complex eukaryote. Proto-eukaryotes have been hypothesized to live in smaller populations than prokaryotes which would have weakened selection for fast growth and thereby allowed genome expansion (Lynch and Conery, 2003). In line with the theory of constructive neutral evolution (Stoltzfus, 1999), the complexity of some molecular machines appears to be redundant (Finnigan et al., 2012; Ba et al., 2017). Generally however, the complex structures and behaviors that arose in eukaryotes are considered to be beneficial.

For the symbiont, gene loss through endosymbiotic gene transfer was supplemented by gene deletion. Once an organism is evolutionarily confined inside a host, Muller’s ratchet can explain genome shrinkage (Nowack et al., 2016). Additionally, adaptive forces may drive genome shrinkage by reducing the cost of genome maintenance and replication. The photosynthetic endosymbiont of *Paulinella chromatophora* which was acquired through a relatively recent primary endosymbiosis event has lost numerous genes that are essential for autonomous growth (Gabr et al., 2022). Similarly, intracellular parasites are hypothesized to have small genomes because the relatively constant host environment does not demand behavioral complexity (McCutcheon and Moran, 2012). Thus, depending on the details of the endosymbiotic relation, both adaptive and non-adaptive forces are likely to play a role in genomic streamlining of endosymbionts.

## Methods

### Internal model of cell-cycle regulation of hosts and symbionts

To study endosymbiosis separate from the metabolic context, we model the autonomous regulation of host and symbiont cell-cycles using our recently published model (Von der Dunk et al., 2022, Fig. 1a, top). In this model, a gene regulatory network arising from stochastic interactions between genes and binding sites on a genome generates cyclic expression dynamics that represent a cell-cycle. Rather than being evaluated with an explicit fitness criterion, cells have to execute the correct cell-cycle stages after which they divide. In particular, the S-stage should be visited sufficiently many times to allow the entire genome to be replicated. Replication speed—i.e. the stretch of genome replicated for one timestep spent in S-stage—is given by the nutrient abundance which depends on preset influx rate into the environment and local population density. Thus replication speed is directly linked to genome size and small genomes are favored because they allow cells to execute faster cell-cycles. Through genome replication, the genome organization also feeds back to the regulatory level as genes are replicated at different times during the cell-cycle, and gene copy numbers influence binding probabilities. We have previously shown that the interaction between genome, gene regulatory network and cellular behavior allows for emergent mechanisms and aids the evolution of complex behaviors (Von der Dunk et al., 2022).

**Figure 1:**
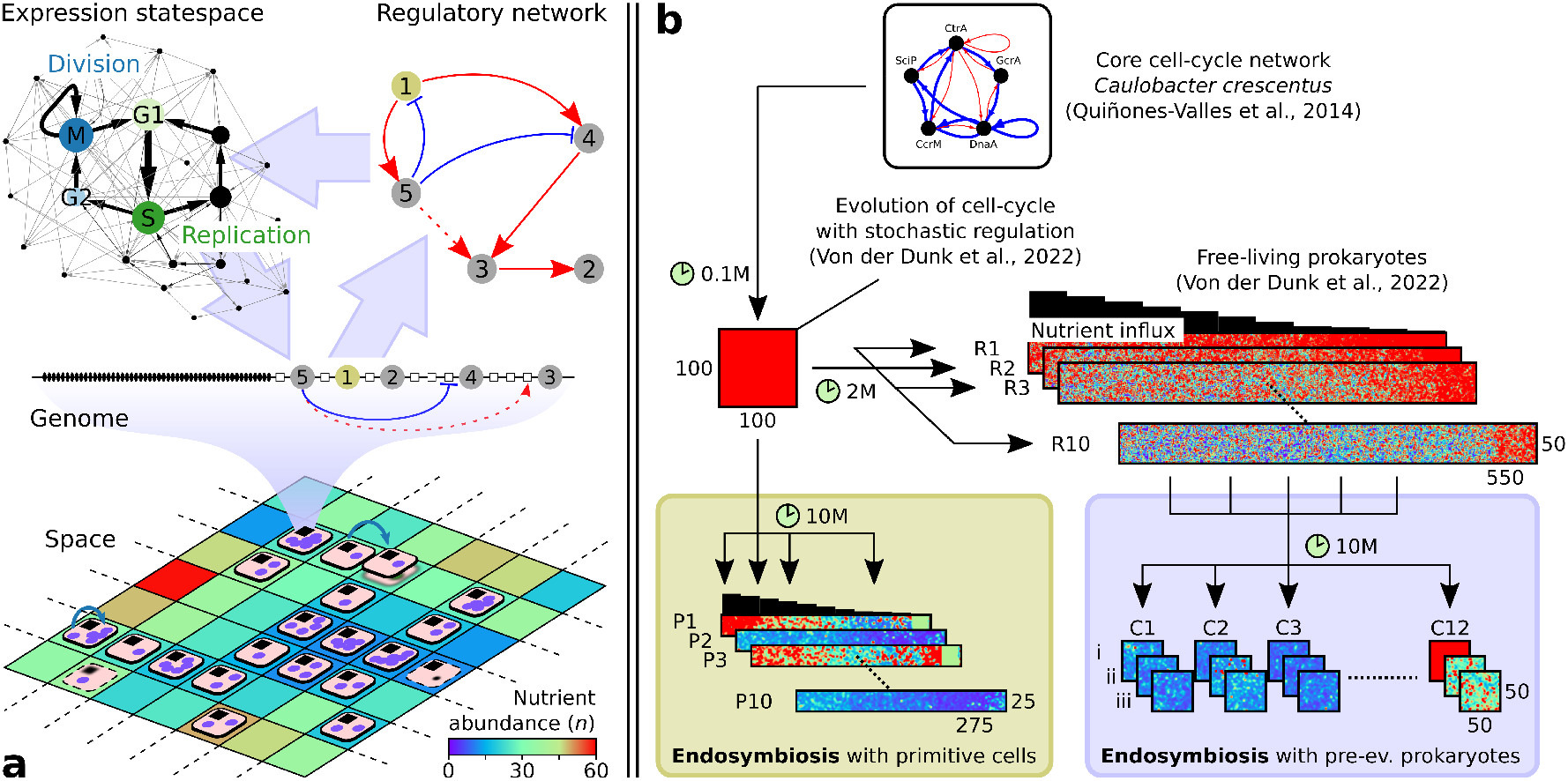
Overview of the model and experimental setup. (a) Host and symbiont each consist of a genome (with discrete beads representing genes and binding sites), gene regulatory network and a cell-cycle. Holobionts (disks) live in space and comprise one host (black square) and one or more symbionts (purple circles) that compete for nutrients. For both host and symbiont, a division event (blue arrow) and a death event (blurred shape) is shown. Lattice site colors depict nutrient abundance for an influx of *n*_*influx*_ = 30. (b) Two evolution experiments with endosymbiosis were performed (green and purple boxes): one starting with primitive cells on a nutrient gradient and one starting with pre-evolved prokaryotes from our previous work (Von der Dunk et al., 2022). Simulation time and grid sizes are shown for each experiment; colors depict nutrient abundance.

### Constructive model of obligate endosymbiosis

Using the above formalism we model endosymbiosis in an individual-based, spatially-embedded system (Fig. 1a, bottom). The simplest way to implement obligate endosymbiosis is to not allow free-living hosts and symbionts. The holobiont—occupying one grid site—thus consists of a single host and one or more symbionts each of which carries its own genome and executes its own cell-cycle as described above. The holobiont dies when the host dies or when the last symbiont dies, which can be due to random death (*d* = 0.001) or due to reaching M-stage prematurely, i.e. without having finished genome replication or without having expressed all preceding cell-cycle stages. The holobiont divides when the host divides by reaching M-stage after a complete cell-cycle. The new holobiont then overgrows one of the neighboring sites, killing the previous occupant if any, and inherits each symbiont of the parental holobiont with a probability of 0.5. Due to stochastic inheritance of symbionts, obligate endosymbiosis forces holobionts to maintain at least two and likely more symbionts at division time to produce viable offspring.

Besides their obligate relationship, hosts and symbionts compete for nutrients in the environment. At each site, nutrients that flux in are equally divided over all hosts and symbionts in the 3-by-3 neighborhood. Through nutrient depletion, high symbiont numbers lead to very slow replication rates which can cause stagnation of the host cell-cycle eventually killing the holobiont. In sum, holobionts require neither too few nor too many symbionts to be both stable (ensuring that offspring receives at least one symbiont) and competitive (ensuring fast growth and preventing host starvation).

Genomes of real organisms encode many genes that are not related to cell-cycle regulation, such as metabolic genes. In our model, each genome carries 50 household genes that are required to perform other tasks for the host or symbiont. To study hypotheses on genome size evolution, we allow these household functions to be encoded by either the host or the symbionts or by both, requiring only that they together encode at least 100 household genes (wherein those of the symbionts are averaged). Through duplication and deletion, household genes can be “transferred” indirectly between host and symbiont, which, along with changes in regulatory repertoire sizes, allows for the evolution of a large host and small symbiont genome or vice versa.

### Eukaryogenesis from primitive or complex FECA

To study the cellular and genomic impact of obligate endosymbiosis, we performed two evolution experiments initialized with holobionts of different complexity which we term primitive and complex FECA, inspired by the First Eukaryotic Common Ancestor (Fig. 1b). For the first experiment, we selected 12 different host–symbiont pairs (C1–12; complex FECA) from the final successful free-living prokaryotes that evolved in Von der Dunk et al. (2022) (see Table S1). For each pair, we ran three technical replicates (i–iii), allowing us to determine how much these initial hosts and symbionts influenced the outcome of evolution. Holobionts adapted quickly and often successfully to the endosymbiotic condition, as hosts and symbionts were already capable of executing complex cell-cycle behavior. Yet the pre-evolved regulatory topologies were not very amenable to changes, so cells turned out to be restricted in their ways of adapting to endosymbiosis as shown below.

For the second experiment, we evolved cell-cycle behavior “from scratch” (P1–10; primitive FECA), i.e. starting with the most primitive cell-cycle network that was also used to initiate evolution in our previous study (Von der Dunk et al. (2022), derived from *Caulobacter crescentus* in Quiñones-Valles et al. (2014)). As in those previous experiments, evolution was carried out on a nutrient gradient where cells could adapt to different nutrient conditions. Since initial regulation was very primitive, cell-cycle adaptation could be accomplished in multiple ways yielding substantial variation in evolutionary trajectories between replicate experiments. Yet adaptation was slow and most populations adapted poorly compared to the evolution experiments with pre-evolved cell-cycle behavior. Nevertheless, we will focus on the replicates starting with a primitive FECA (P1–10) to disentangle how cell behavior has adapted to endosymbiosis, and then turn our focus to the replicates starting with a complex FECA (C1–12) to understand how genome size evolution interacts with endosymbiosis.

## Results

### Holobionts adapt to nutrient gradient

Fig. 2a shows the adaptive process of the ten replicates starting with primitive hosts and symbionts (P1–10). In three replicates, the small initial population was unable to improve cell-cycle behavior and stabilize the holobiont, resulting in early extinction before *t* = 10^6^. Yet in the seven remaining replicates, holobionts managed to adapt to limited nutrient conditions and the endosymbiotic lifestyle, reaching final population sizes ranging from about 28% (P6) to 63% (P9) of the grid. Thus, the experiment has successfully generated diverse evolutionary trajectories emanating from the same origin and experiencing the same environmental conditions.

**Figure 2:**
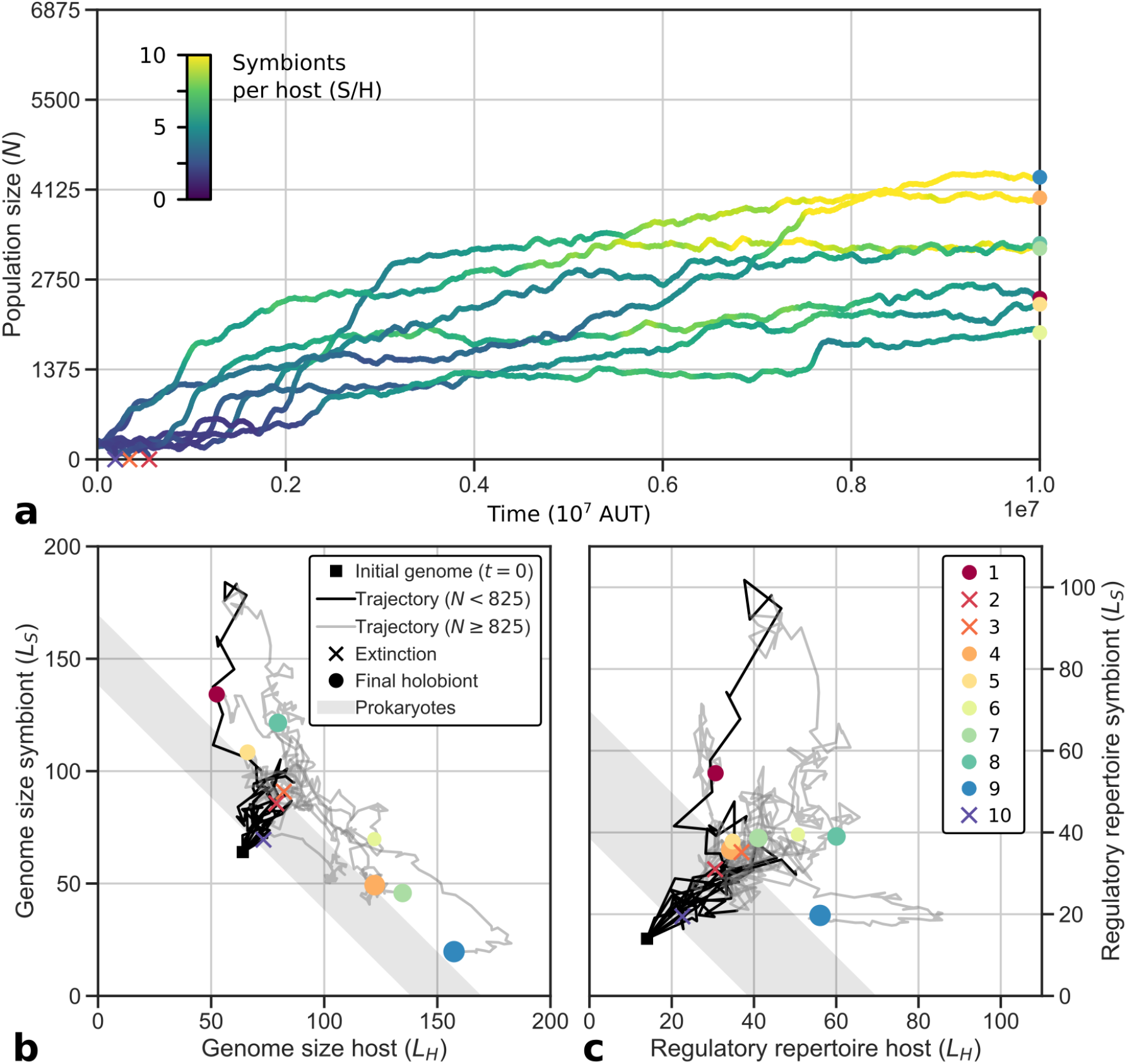
Primitive holobionts adapt to nutrient gradient. (a) Population size increases, coinciding with increases in symbiont numbers. (b) Genomes expand early, and later genome size symmetry breaks between host and symbiont. (c) Regulatory repertoires expand further than free-living cells for all hosts and symbionts that survive. Maximum population size is given by the grid size: 25 *×* 275 = 6875. Adaptation generally occurs before *t* = 10^7^ (see Fig. S1 for trajectories until *t* = 2·10^7^). In panels b and c, the size of the markers is scaled by final population size.

Populations grow by expanding their range on the nutrient gradient and by increasing their density. In the most successful replicate (P9), we can distinguish two adaptive phases (*t* ≈ 10^6^ and 5·10^6^ < *t* < 8·10^6^) when the population expands its range to poorer conditions and simultaneously increases its density in rich conditions (Fig. 3). In general, we find that adaptation occurs in relatively short phases interspersing long periods of stasis (Fig. 2a, viz. Gould and Eldredge, 1972), which is similar to what we found in our previous model that represents free-living prokaryotes (Von der Dunk et al., 2022).

**Figure 3:**
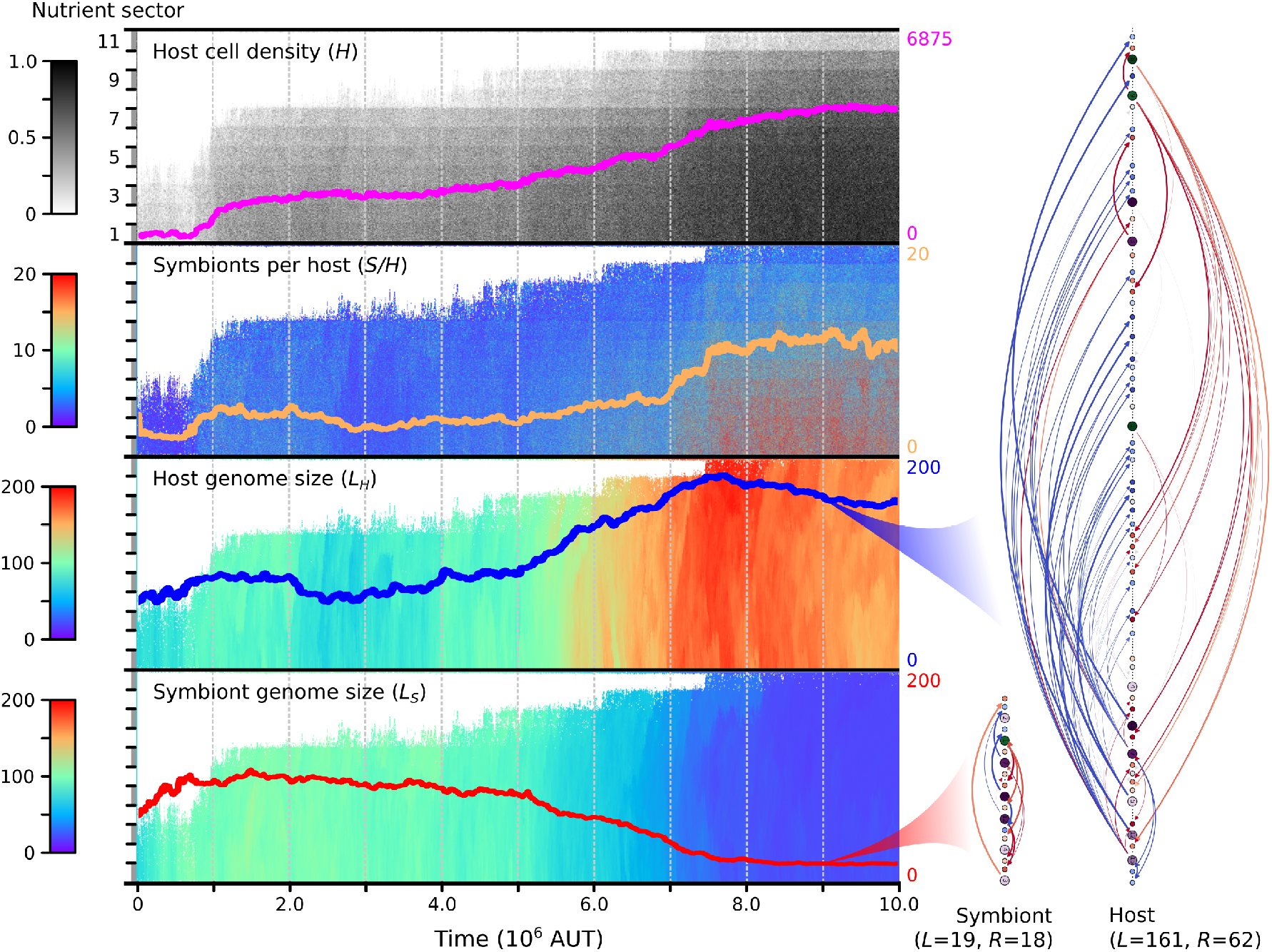
Evolutionary dynamics of the most successful replicate with primitive FECA, P9. Each panel shows a different variable in spacetime. The overlaid lines represent summaries across the entire gradient (for the top panel, overlaid line shows total population size rather than local cell density, in correspondence with Fig. 2a). On the right, genomes of the host and symbiont in the ancestor at *t* = 9 · 10^6^ are depicted from top to bottom, along with the regulatory interactions (*L* denotes total genome size, *R* regulatory repertoire size). For an example of a different evolutionary trajectory, see Fig. S2.

### Genome expansion and adaptation

In all replicates, host and symbiont genomes expand rapidly early in evolution (Fig. 2b,c). Large genomes are *prima facie* unfavorable because they take long to replicate, forcing cells to execute slow cell-cycles. Interestingly, the rapid expansion occurs when holobionts are still primitive and unfit, at population sizes below 12% of the grid (*N <* 825). The three replicates that went extinct experienced genome expansion during their entire existence.

Early genome expansion thus apparently results from a lack of selection. At the same time, expanding the regulatory repertoire allows hosts and symbionts to invent more complex cell-cycle behavior and thereby underlies the first substantial adaptations in the seven replicates that survive (e.g. the adaptation seen in P9 around *t* = 10^6^ in Fig. 3). These adaptations lead to an increase in population size, after which selection intensifies and genome expansion slows down.

At the end of the experiment, the regulatory repertoires of hosts and symbionts have surpassed what we observed for free-living prokaryotes (the grey band in Fig. 2c; cf. Von der Dunk et al., 2022). Endosymbiosis imposes constraints on host and symbiont that are absent in free-living cells and which weaken selection for genomic streamlining. Thus, the endosymbiotic condition contributes to genome expansion and potentially explains some of the complexity that emerged during eukaryogenesis.

### Concurrent expansion and shrinkage of host and symbiont genomes

On long timescales, the genome sizes of host and symbiont diverge: four replicates evolve towards large host and small symbiont genomes and three replicates evolve towards large symbiont and small host genomes (Fig. 2b). The first of these two scenarios, which resembles the outcome of eukaryogenesis, is also more common in the evolution experiment starting with complex FECA (C1–12). Specifically, the replicates that are initialized with identical host and symbiont (C1–4) always evolve to have a somewhat larger host than symbiont genome. Moreover, in both evolution experiments (P1–10 and C1–12), the replicates that evolved a larger host than symbiont genome reached larger population size (see Fig. S3). In P9, the dramatic symmetry breaking in genome size that takes place from around *t* = 4 · 10^6^ coincides with an increase in population size revealing adaptation until *t* = 8 · 10^6^. Thus, successful endosymbiosis leads to a complex host and a simple symbiont, similar to the outcome of eukaryogenesis.

### Evolution of symbiont numbers

Symbiont numbers constitute an important degree of freedom of our model that is subject to enormous variation arising at different levels. For an individual holobiont, there is already an associated probability distribution of symbiont numbers due to inherent stochasticity in regulatory dynamics and due to changes in host and symbiont cell-cycle behavior as a result of variation in nutrient conditions. For the entire population, mutations create differences in these probability distributions between individual holobionts which allows them to evolve over time. Below, we describe how variation in symbiont numbers across time and space has contributed to holobiont adaptation, starting at the population-level, then zooming in to individual holobionts, and finally focusing on the details of host and symbiont cell-cycle behavior within a single holobiont.

During the evolution experiment initialized with primitive FECA, the average number of symbionts per host in the population increases from 2.5*−*2.8 to 4.3*−*10.2 across replicates (Fig. 2a). The increase in symbiont numbers is directly linked to holobiont adaptation. In P9 for instance, both adaptive phases that we identified before, are accompanied by marked increases in symbiont numbers (Fig. 3). More generally, final population size correlates well with symbiont number across replicates (*r* = 0.68; Fig. S4). Thus, symbiont numbers play an important role in adaptation towards a successful endosymbiotic lifestyle. On top of the increased average symbiont number at the end of replicate P9, we find that symbiont numbers have diversified across the nutrient gradient (this is the case for about half of the replicates; other populations evolved low symbiont numbers and little diversity across the gradient, see e.g. Fig. S2).

### Holobionts adapt through r- or K-strategy

To understand the observed evolutionary patterns in genome size and symbiont number, we now study adaptive behavior at the level of individual holobionts, which will subsequently be broken down further into the underlying host and symbiont behaviors. First, to discern the strategies by which holobionts have adapted, we studied clonal population dynamics of four ancestors extracted at different timepoints along the evolutionary trajectory of P9 under the nutrient conditions of each sector of the gradient (see “Characterizing holobionts” in Supplementary Material; Fig. 4). As expected, these ancestors broadly recapitulate the adaptation that was observed in the full population, i.e. increases in host cell density and symbiont numbers (Fig 4, top panels). In addition, holobiont profiling reveals two alternative strategies (Fig. 4). The early ancestors at *t* = 0 and *t* = 3 · 10^6^ maintain very few symbionts, yielding unstable holobionts and low cell density. As a consequence, nutrients are not depleted much and holobionts grow (i.e. invade) fast in those conditions where they are viable (*n*_*influx*_ < 50 and *n*_*influx*_ < 10, respectively). Conversely, the more recent ancestors at *t* = 6 · 10^6^ and *t* = 9 · 10^6^ maintain many symbionts, yielding stable holobionts and high cell density. Yet nutrients are depleted to low levels and holobionts grow slow. Thus, through evolution of symbiont numbers, holobionts have access to an r-strategy which optimizes growth (*r*) and a K-strategy which optimizes carrying capacity (*K*). Early in evolution, when empty space and nutrients are abundant, selection for fast invasion favors the r-strategy, whereas later, as empty space and nutrients become limited due to improved cell-cycle behavior, the K-strategy is more advantageous. Nevertheless, some populations settle on an r-strategy (e.g. P8, C10 and C12; see also “Genome size evolution directs holobiont adaptation”), highlighting the role of historical contingency and chance processes in evolution.

**Figure 4:**
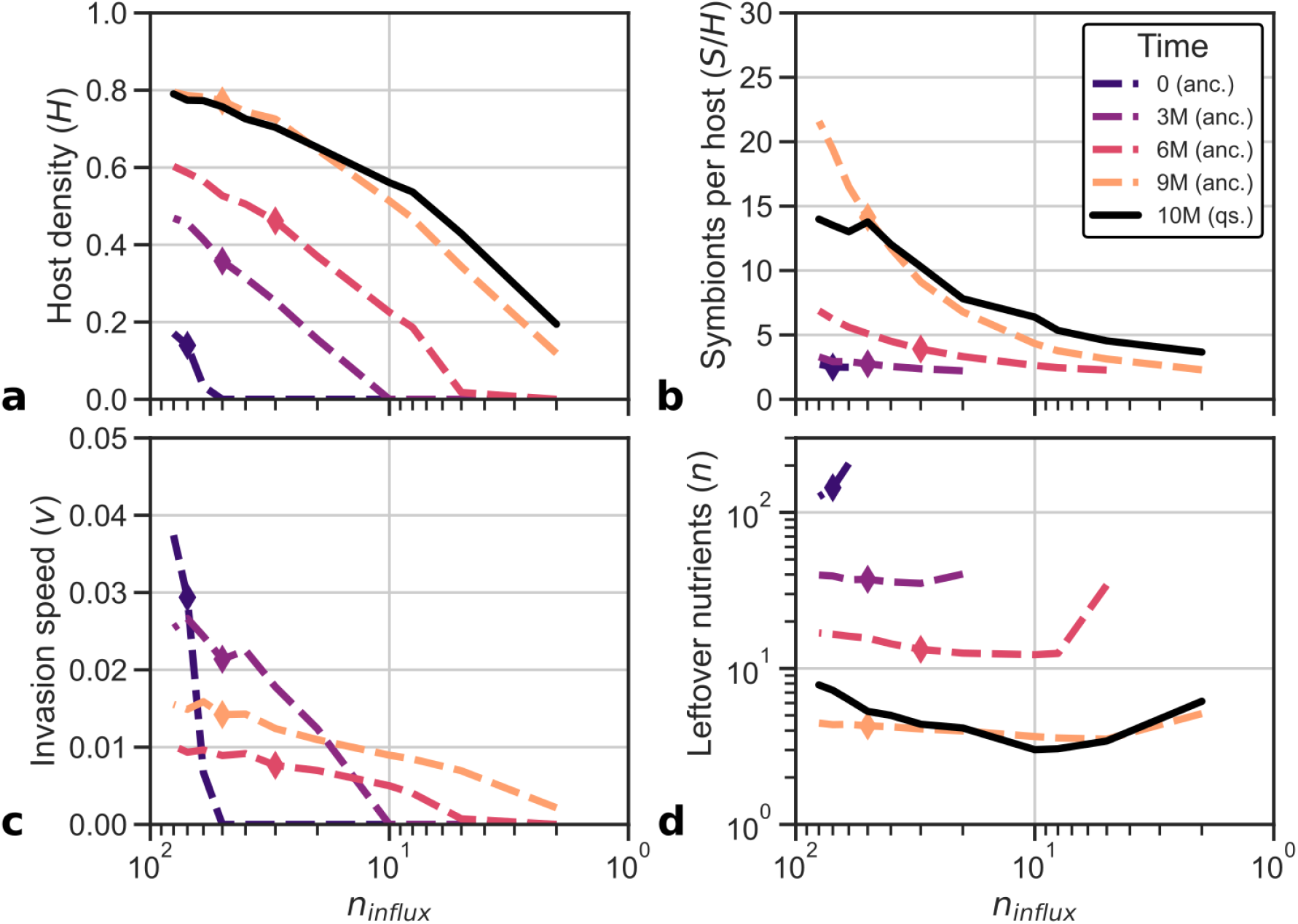
An r- and K-strategy have emerged along the ancestry of P9. Over time, cell density (a) and symbiont number (b) have increased, but growth measured as invasion speed (c) and nutrient availability (d) have decreased. The final population resembles the most recent ancestor but strains at high and low nutrient influxes have adapted to those specific environments. Diamonds depict the conditions in which the ancestor lived.

### Individual- and population-level regulation of symbiont numbers

Surprisingly, the previously observed variation in symbiont numbers across the nutrient gradient at the end of replicate P9 (Fig. 3) is for a large part explained by individual regulation of symbiont numbers with nutrient conditions as observed in the ancestor profiles (Fig. 4b). When the holobiont encounters poor nutrient conditions, it maintains few symbionts, and when it encounters rich nutrient conditions, it maintains many symbionts. This emergent control ensures that there are always sufficient nutrients available for the host to finish genome replication (as seen in Fig. 4c) and that resources are only diverted to symbionts to increase holobiont stability when conditions allow it. Thus, the holobiont has evolved a mechanism to successfully navigate the large fluctuations in nutrient levels resulting from local variation in host cell density and symbiont numbers.

The full quasi-species at *t* = 10^7^ diverges only slightly from the individual symbiont control observed in the ancestor at *t* = 9 · 10^6^. In environments that are poorer than the native environment of the ancestor (*n*_*influx*_ = 50), strains of the quasi-species have adapted by boosting average symbiont numbers relative to the ancestor (e.g. from 2.3 to 3.7 at *n*_*influx*_ = 2), enhancing holobiont stability and carrying capacity, and reducing growth. Conversely, in richer environments, strains have adapted by reducing symbiont numbers relative to the ancestor (e.g. from 21.6 to 14.0 at *n*_*influx*_ = 80), promoting growth while barely reducing carrying capacity—since stability is already almost guaranteed with more than ten symbionts per host.

### Emergent symbiont control through implicit cell-cycle coordination

To understand how the adaptations to endosymbiosis at the holobiont level come about— i.e. r- and K-strategies and symbiont control—we analyse the cell-cycle behavior of host and symbiont independently (see “Tracking cell-cycle behavior” in Supplementary Material). By plotting their growth curves as a function of nutrient condition (see “Calculating growth rate” in Supplementary Material), we obtain a phase diagram for the holobiont (Fig. 5, left panels). For any particular nutrient condition, the relative positions of the growth curves reveal whether symbiont number will increase (*r*_*S*_ > *r*_*H*_) or decrease (*r*_*H*_ > *r*_*S*_) with time. The change in symbiont number translates to a shift on the x-axis: an increase in symbiont number corresponds to a decrease in nutrient abundance (shift to the right), and a decrease in symbiont number corresponds to an increase in nutrient abundance (shift to the left).

**Figure 5:**
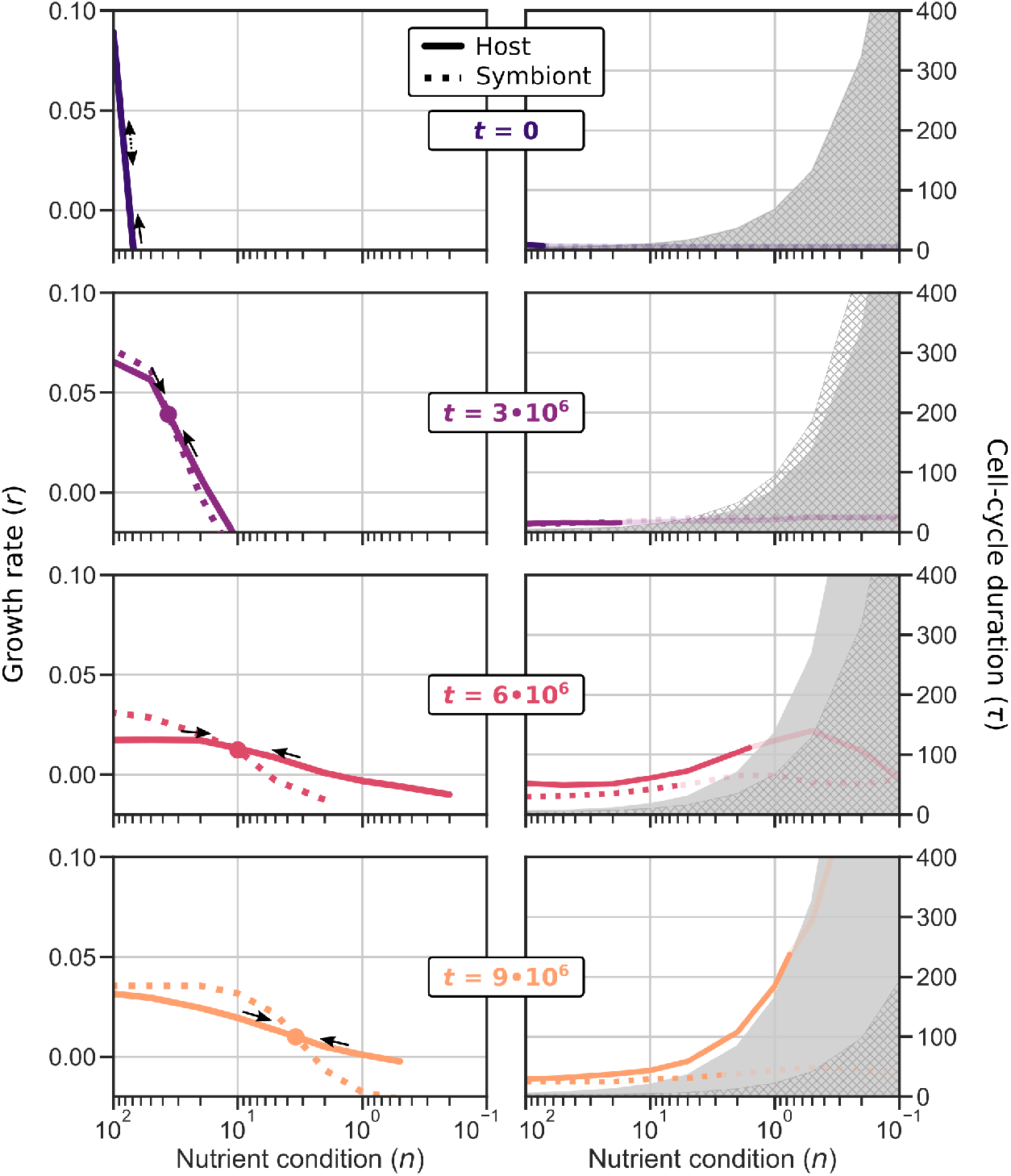
Host–symbiont cell-cycle coordination evolves, first through differential cell-cycle timing, later through niche differentiation whereby the host becomes a generalist and the symbiont a specialist. Each row shows the growth rates (left panels) and cell-cycle durations (right panels) of an ancestor of the P9 replicate. In contrast to Fig. 4, the x-axis here depicts fixed nutrient condition where host and symbiont cell-cycle dynamics are assessed independently. The grey areas in the cell-cycle duration panels mark the minimum possible duration of a successfull cell-cycle (with enough time for replication) given the genome size of the host (filled) and symbiont (hatch). The cell-cycle duration is only drawn in bold for the range where *ρ* > 0.5, which roughly coincides with *r* > 0

The phase diagrams depicting growth of ancestral hosts and symbionts reveal how holobionts achieved symbiont control during evolution. Initially, symbiont dynamics are unstable: even with equal host and symbiont growth rate across nutrient conditions, stochasticity causes symbiont number to drift up or down, resulting in holobiont death due to symbiont loss or host starvation (i.e. *r*_*H*_ < 0 for *n* ≲ 70 for the initial holobiont). Later holobionts (*t ≥* 3 · 10^6^) managed to stabilize symbiont dynamics through differentiation of host and symbiont growth. Perturbations from the stable equilibrium (i.e. the intersection of *r*_*H*_ and *r*_*S*_) are compensated by subsequent changes in relative symbiont and host growth rate that push the system back to this equilibrium. The evolved stable equilibrium also explains previously observed holobiont behavior: host–symbiont growth dynamics equilibrate at the same nutrient level regardless of nutrient influx (Fig. 4d), which means that with greater nutrient influx more symbionts can be sustained before nutrients are depleted to that equilibrium level (Fig. 4b).

Given that we understand how holobionts control symbiont numbers, we can clearly observe how cells evolved higher symbiont numbers over time (Fig. 5). Between *t* = 3 · 10^6^ and *t* = 9 · 10^6^, the position of the stable growth point moves from high nutrient abundance (few symbionts) and high growth rate (r-strategy) to low nutrient abundance (many symbionts) and low growth rate (K-strategy). Underlying this transition from r-to K-strategy are changes in the cell-cycle behavior. First, the host growth curve is flattened when the host has evolved a slower cell-cycle and increased survival in poor conditions (e.g. *t* = 6 · 10^6^). Later (*t* = 9 · 10^6^), host and symbiont diverge dramatically in genome size and differentiate into a generalist and specialist, respectively, referring to their ability (and inability) to tune cell-cycle behavior to nutrient conditions (Von der Dunk et al., 2022). The invention of generalism allows the holobiont to maintain more symbionts in equilibrium increasing stability while also promoting faster growth when provided with more nutrients at the invasion front (i.e. see Fig. 4c).

### Cell-cycle coordination and genome size evolution

The characterization of cell-cycle coordination between host and symbiont allows us to revisit emergent genome size asymmetry, i.e. the appearance of complex hosts with simple symbionts or vice versa (Fig. 2b). For this, we turn our focus to the evolution replicates starting with a complex FECA (C1–12). It turns out that when a prokaryote with a more efficient cell-cycle (i.e. R8 or R9)—quantified as 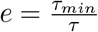 (see Supplementary Material)— is matched with a prokaryote with a less efficient cell-cycle (e.g. R2 or R3), the former evolves a large genome and the latter a small genome, irrespective of which prokaryote is the host and which the symbiont. Across both evolution experiments, asymmetry in cell-cycle efficiency explains 92% of the variation in genome size asymmetry (Fig. 6).

**Figure 6:**
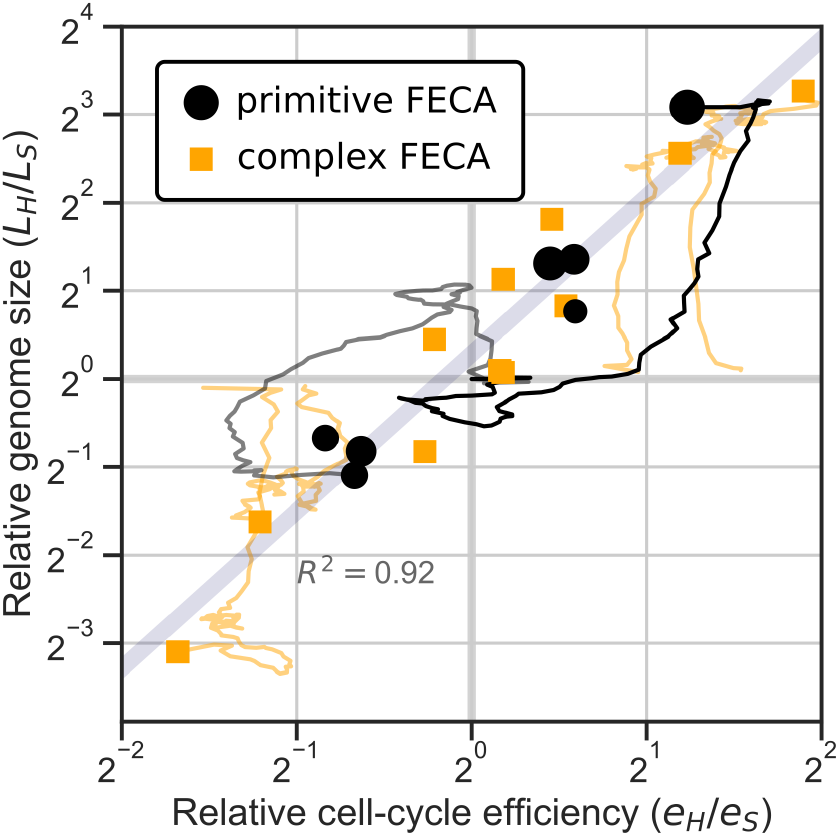
Asymmetry in genome size correlates strongly with asymmetry in cell-cycle efficiency between host and symbiont. Along the ancestral lineage, genome size and cell-cycle efficiency are measured (see “Calculating cell-cycle efficiency” in Supplementary Material). Trajectories are shown for the experiments where the largest genome size asymmetry evolved (P1, P9 and C9–12); the trajectory of P9 is in bold. One technical replicate is shown for each of the replicates initialized with complex FECA. For the experiment initialized with primitive FECA, the size of the markers is scaled by final population size.

The strong correlation between cell-cycle efficiency and genome size in host–symbiont pairs is the result of selection for cell-cycle coordination. To balance growth between host and symbiont, selection forces the more efficient partner to slow down its cell-cycle to the speed of the less efficient partner. The added cell-cycle time is utilized by the more efficient partner to replicate more household genes, relieving the less efficient partner of this burden. Thus, genome size asymmetry compensates for differences in cell-cycle efficiency between host and symbiont, allowing for faster overall growth of the holobiont. This mechanism applies when asymmetry in cell-cycle efficiency is ingrained in the holobiont from the start, as in the evolution experiment with complex FECA, but also when the asymmetry arises spontaneously, as in the evolution replicates with primitive FECA where, by chance, either the host or symbiont acquires more efficient cell-cycle regulation.

A similar adaptive mechanism explains why hosts evolve larger genomes than symbionts when they start out identical (C1–4). Hosts generally evolve slower cell-cycles than symbionts in order to stabilize symbiont dynamics (i.e. to obtain the stable dynamic equilibrium, Fig. 5), so they can spend more time replicating household genes and partly relieve the symbiont.

As previously mentioned, non-adaptive forces play a large role in genome expansion and genome size early in evolution. Yet the final asymmetry in genome size is dominated by adaptive forces compensating differences in cell-cycle efficiency between host and symbiont (Fig. 6). When household genes cannot be shared between host and symbiont, a small but consistent asymmetry remains (Fig. S5), indicating that a host or symbiont with greater replication capacity due to a slower cell-cycle experiences more genome expansion than its partner even when there is no compensatory advantage by concurrent streamlining of that partner genome. In short, large genomes are less costly when a slow cell-cycle is favored by other constraints.

### Genome size evolution directs holobiont adaptation

Genome size evolution is not just the byproduct of selection for cell-cycle coordination. In several replicates, a marked switch in holobiont behavior evolves after extensive symmetry breaking in genome size. In two replicates, C11 and C12, the evolutionary trajectories intersect in phenotypic space (Fig. 7): C11 evolves from low cell density and low symbiont number (r-strategy) to high cell density and high symbiont number (K-strategy), and C12 follows this path in reverse. In both cases, competition experiments between populations at different times show that each evolutionary path is entirely adaptive from before until after the switch, and this is supported by the congruence between technical replicates.

**Figure 7:**
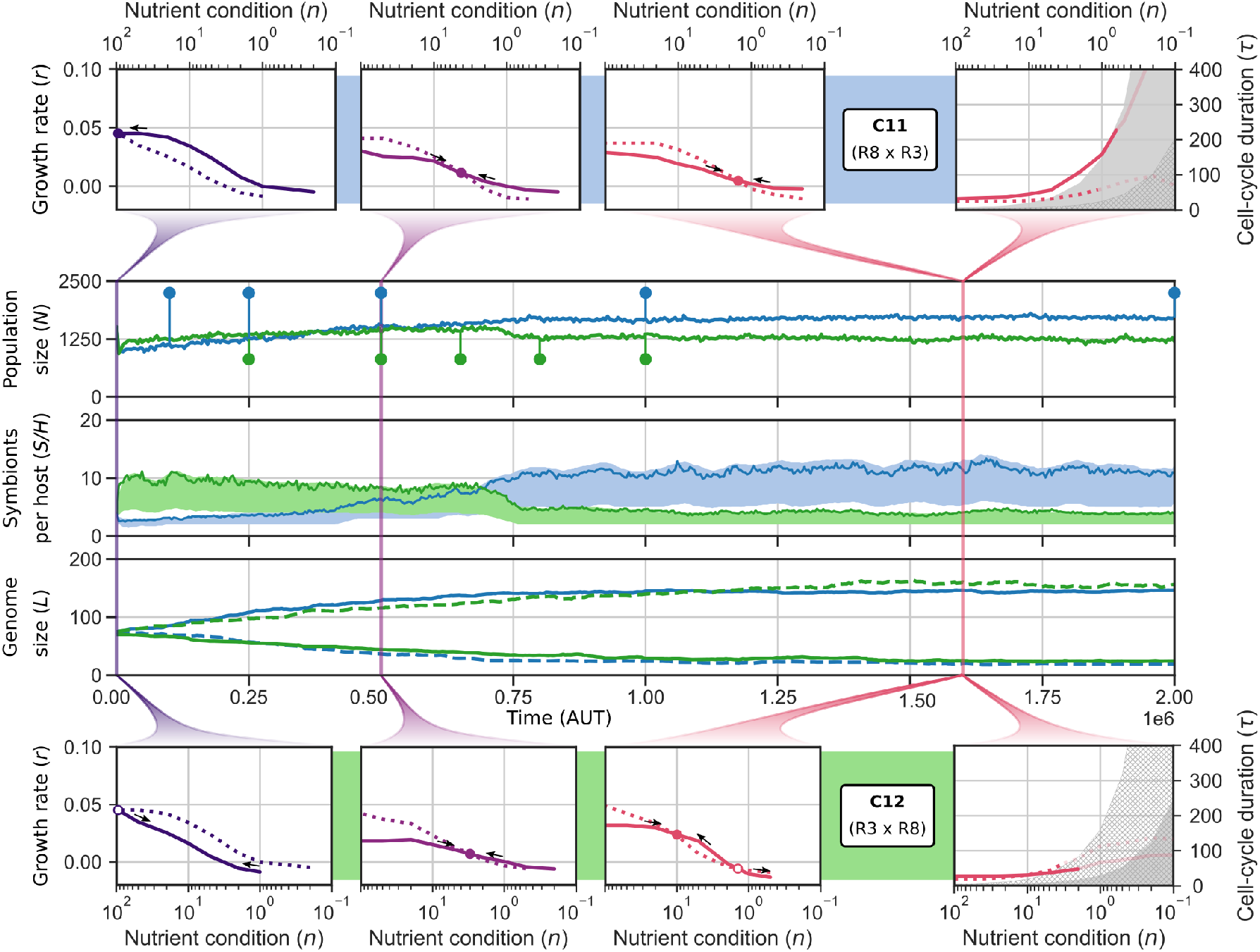
Strategy switching through symmetry breaking in genome size in C11 and C12. These two populations follow opposite trajectories, and intersect in several aspects: population size, symbiont number and growth dynamics. For each replicate, competition experiments between populations from different timepoints (circles in population size panel) revealed unequivocal fitness increase across the strategy switch. The cell-cycle behavior (top and bottom right-most panels) show that cell-cycle behavior is very different between C11 and C12, explaining why holobionts traverse opposite evolutionary trajectories. For symbionts per host (middle panel), the shaded area spans the 33% and 67% quantiles; the lines show the means (distributions are skewed).

The critical difference between C11 and C12 which explains why seemingly identical holobionts (around *t* = 0.5 · 10^6^) continue to evolve in diametrically opposite directions, is the distinct cell-cycle behavior that underlies growth coordination. In C11, holobionts are formed by a generalist host (R8) and a specialist symbiont (R3) resembling P9, whereas in C12, holobionts are made up of a specialist host (R3) and a generalist symbiont (R8; as clearly seen in the top and bottom right-most panels in Fig. 7). These contrasting host–symbiont behaviors can generate very similar growth dynamics (i.e. phase diagram at *t* = 0.5 · 10^6^), but ultimately give rise to different constraints and different opportunities for adaptation. Genome size asymmetry is the key factor that opens up the different adaptive routes: when we disallow sharing of household genes between host and symbiont, genome size asymmetry is less extreme and the strategy switches do not occur.

On the long term, the trajectories of C11 and C12 are not equally adaptive. The final population of C11 (at *t* = 10^7^) outcompetes that of C12, reiterating that holobionts with large host and small symbiont genomes—i.e. those resembling eukaryotes—are fitter.

More specifically, the r-strategy yields small symbiont populations with very large genomes, which weakens selection and reduces competitiveness late in the experiment (around *t* = 8 · 10^6^).

## Discussion

### Endosymbiosis begets complexity

We forced different combinations of cells with their own genome and gene regulatory network into an obligate endosymbiotic relationship. In the vast majority of our experimental replicates, holobionts evolved implicit cell-cycle coordination through differential growth behavior on the same resource. This shows that no complex cellular traits are in principle required to control symbionts other than the ability of host and symbiont to adapt to different nutrient conditions, i.e. no intracellular communication via protein targeting is required and no advanced cell-cycle control by the host.

Nevertheless, not all cells become equally successful hosts or symbionts. Pre-evolved cells which already performed complex cell-cycle regulation adapted much faster to endosymbiosis and reached greater population sizes than primitive cells. As in free-living cells, the efficiency of cell-cycle regulation is an important factor in successful holobiont adaptation. In addition, generalist behavior is preferred for hosts, allowing them to deal with both high and low nutrient supplies resulting from fluctuations in symbiont numbers. For symbionts, relatively simple specialist behavior as observed in P9 appears sufficient for successful holobionts. In nature, endosymbionts experience a relatively constant environment inside their host which has been hypothesized to drive genomic streamlining (McCutcheon and Moran, 2012; Gabr et al., 2022). In our model, nutrient homeostasis is accomplished through indirect control of symbiont numbers and by adaptation of the host to the remaining fluctuations, which also allows the symbiont to specialize on specific nutrient conditions and streamline its genome.

### Genome size evolution

During eukaryogenesis, genome size increased immensely (Vosseberg et al., 2021). In our model, the total genome size of the holobiont also increased with respect to free-living cells where genome expansion was limited but necessary for functional adaptation (Von der Dunk et al., 2022). Moroever, under the high mutation rates of the “prokaryotic” regime, holobionts experienced even more genome expansion and went extinct frequently, indicating that the endosymbiotic condition weakens selection for genomic streamlining. These outcomes are in line with the non-adaptive account of genome expansion proposed by Lynch (Lynch and Conery, 2003). When an organism has more tasks to perform than replicating as fast as possible, selection for small genomes to speed up replication is weakened.

Another striking pattern in the outcome of eukaryogenesis is the asymmetry in genome size between the host (nuclear) and symbiont (mitochondrial) genomes (Roger et al., 2017). Most of the proposed drivers of this asymmetry—such as increased substitution rate in the symbiont due to lack of recombination and ROS production (Allen and Raven, 1996; Martin and Herrmann, 1998) or advantageous host regulatory control (Nowack, 2014)—are not incorporated into our model, yet asymmetry still evolves. In our model, unequal replication efficiency drives genome size asymmetry. Apart from the outcome of eukaryogenesis, i.e. large host and small symbiont genome, this mechanism also yields the opposite outcome, i.e. small host and large symbiont genome, which has not been observed in any endosymbiosis in nature. Still, the eukaryogenesis-like outcome is favored because on short timescales, cell-cycle coordination selects for a slow host, yielding asymmetric constraints on genome size, and on long timescales, holobionts with large host and small symbiont genomes evolve the highest carrying capacity (K-strategy) outcompeting all other holobionts.

Since all extant endosymbiotic relationships involve a host that is already complex, host control over symbiont numbers has not yet been addressed as a potential hurdle for eukaryogenesis. Yet for eukaryogenesis we do not know whether targeted protein transport through a complex endomembrane system was already possible. If host control over symbiont numbers evolved later during eukaryogenesis, the number of symbionts per holobiont would initially have been a key emergent property of host and symbiont growth dynamics. As we show in our relatively simple model where symbiont numbers impact holobiont behavior and interact with the environment, multiple holobiont strategies are possible. The evolved r- and K-strategies are tightly linked to genome size evolution. In the K-strategy, symbionts exist in large populations and experience strong selection for genomic streamlining and fast growth. In contrast, in the r-strategy, symbionts evolve large genomes and exist in small populations, which can lead to collapse of the population much later on (e.g. in C12 around *t* = 8 · 10^6^, far outside the range shown in Fig. 7).

A surprising result from this study is that effective cell-cycle coordination between host and symbiont in obligate endosymbiosis can evolve without explicit communication. This raises the question why intracellular communication and host cell-cycle control—e.g. through protein targeting and other signaling pathways—evolved during eukaryogenesis. Strikingly, we have shown that the evolution of large eukaryotic genomes is explained as a consequence of both adaptive forces—selection for cell-cycle coordination—and non-adaptive forces—reduced selection for fast growth—and that the most successful evolution replicates mimic eukaryogenesis. Our results underscore the power of modeling approaches for understanding complex evolutionary processes such as eukaryogenesis and the mitochondrial endosymbiosis.

## Acknowledgements

The authors gratefully acknowledge the help of Jan Kees van Amerongen for running the local computer cluster. This work is part of the research programme VICI with project number 016.160.638, which is financed by the Netherlands Organisation for Scientific Research (NWO).

