## Supplementary Material for "Obligate Endosymbiosis Explains Genome Expansion During Eukaryogenesis"

November 15, 2022

#### Supplementary Methods

##### Characterizing holobionts

To study how cell-cycles of host and symbiont adapted to endosymbiosis, we profiled holobionts in the ancestral lineage of P9. For this, we performed small experiments with clonal populations of a single holobiont type (without mutation) on every sector of the gradient. First, propagation across empty space is measured, followed by bulk measures of cell density, symbionts per host and leftover nutrients. We also measured 3 of the 4 observables directly in the gradient at the end of replicate P9 ($t = 10^7$), to see if and how diversity in the quasi-species has improved adaptation to the gradient relative to the ancestral holobiont at $t = 9 \cdot 10^6$.

##### Tracking cell-cycle behavior

We performed experiments where individual hosts and symbionts execute their cell-cycle behavior for $10^4$ AUT under different fixed nutrient conditions (as described in Von der Dunk et al., 2022). During the experiment, we tracked the average cell-cycle duration $\tau$, the fraction of successful cell-cycles $\rho$, and calculated the instantaneous growth rate as $r = \frac{\rho}{\tau}\left(1 - \frac{1}{R_0}\right)$ where $R_0 = \frac{\rho}{\delta\tau+1-\rho}$ (based on an ODE description of the model, see "Calculating growth rate" and Von der Dunk et al., 2022).

#### Calculating growth rate

In the Supplementary Material of our previous study (Von der Dunk et al., 2022), we presented a simple ODE model that describes phenomenologically a cell-cycle under constant nutrient conditions.

$$\frac{dN}{dt} = \frac{\rho}{\tau}N(1-N) - \frac{1-\rho}{\tau}N - \delta N \tag{1}$$

From the cell-cycle parameters $\tau$ and $\rho$ (which are measured in single-cell simulation runs), we derived $R_0$ as:

$$R_0 = \frac{\rho}{\delta\tau + 1 - \rho} \tag{2}$$

Here we derive the instantaneous growth rate $r$ allowing us to compare the effective growth of the symbiont relative to the host across nutrient conditions. Similar to $R_0$, the instantaneous growth rate is defined in optimal conditions (i.e. $N \to 0$), as the per capita growth rate:

$$r = \frac{\rho}{\tau}(1-N) - \frac{1-\rho}{\tau} - \delta \tag{3}$$

$$r = \frac{2\rho - 1 - \delta\tau}{\tau} \tag{4}$$

$$r = \frac{\rho}{\tau}\left(1 - \frac{1}{R_0}\right) \tag{5}$$

#### Calculating cell-cycle efficiency

The fastest possible cell-cycle for a given fixed nutrient abundance $n$ and genome size $L$ is $\tau_{min} = 3 + L/n$. The duration of an actual cell-cycle relative to this fastest possible cell-cycle gives the efficiency $e = \frac{\tau_{min}}{\tau} = \frac{3+L/n}{\tau}$. We average efficiencies obtained under conditions $n \in \{100, 50, 20, 10, 5, 2, 1\}$ from the single-cell experiments ("Tracking cell-cycle behavior" in Supplementary Methods) to arrive at a single value for an individual.

### Supplementary Tables and Figures

Table S1: Important characteristics of pre-evolved free-living prokaryotes, i.e. cell-cycle efficiency (see Fig. 6) and generalist capacity (plasticity measured as the log-difference between cell-cycle duration at $n = 0.1$ and $n = 100$).

| Strain | Efficiency ($e$) | Plasticity ($\sigma$) |
|---|---|---|
| R8 | 0.476 | 0.965 |
| R9 | 0.402 | 0.866 |
| R2 | 0.307 | -0.355 |
| R3 | 0.160 | 0.497 |

Table S2: Host–symbiont pairs used to initialise evolution experiments (see Table S1 for characteristics of prokaryotes). C1–4 start with identical host and symbiont; C5–6 have different host and symbiont but with very similar phenotypic behavior, i.e. R8 and R9 are both efficient generalists; C7–12 are asymmetric.

| Holobiont | Host | Symbiont |
|---|---|---|
| C1 | R8 | R8 |
| C2 | R9 | R9 |
| C3 | R2 | R2 |
| C4 | R3 | R3 |
| C5 | R8 | R9 |
| C6 | R9 | R8 |
| C7 | R8 | R2 |
| C8 | R2 | R8 |
| C9 | R3 | R2 |
| C10 | R2 | R3 |
| C11 | R8 | R3 |
| C12 | R3 | R8 |

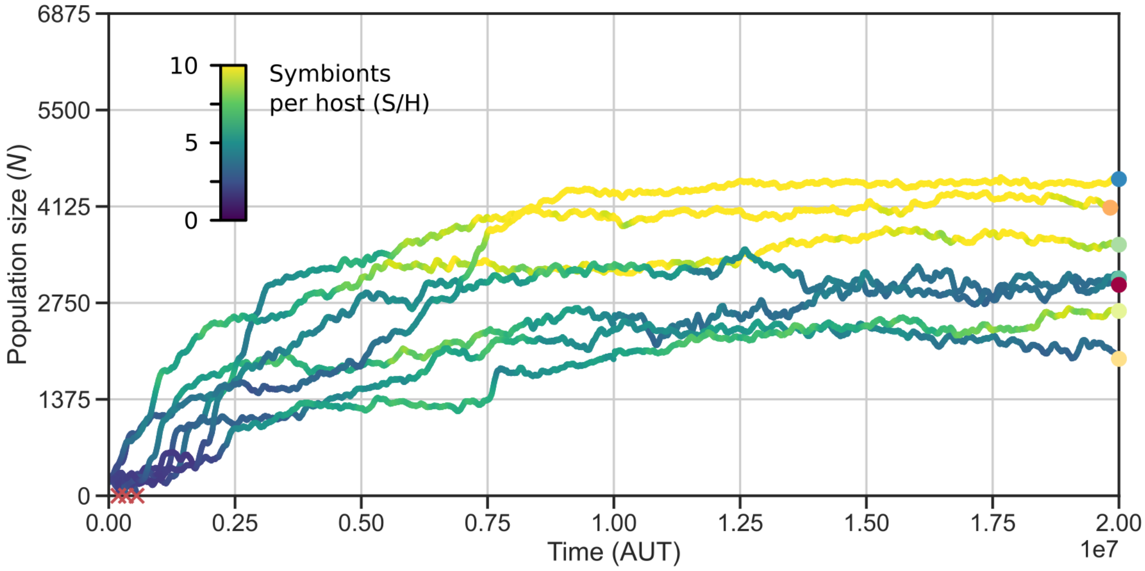


Figure S1: Evolutionary trajectories of the replicates with primitive FECA, continued until $t = 2 \cdot 10^7$, showing that adaptation saturates after $t = 10^7$ (cf. Fig. 2a).

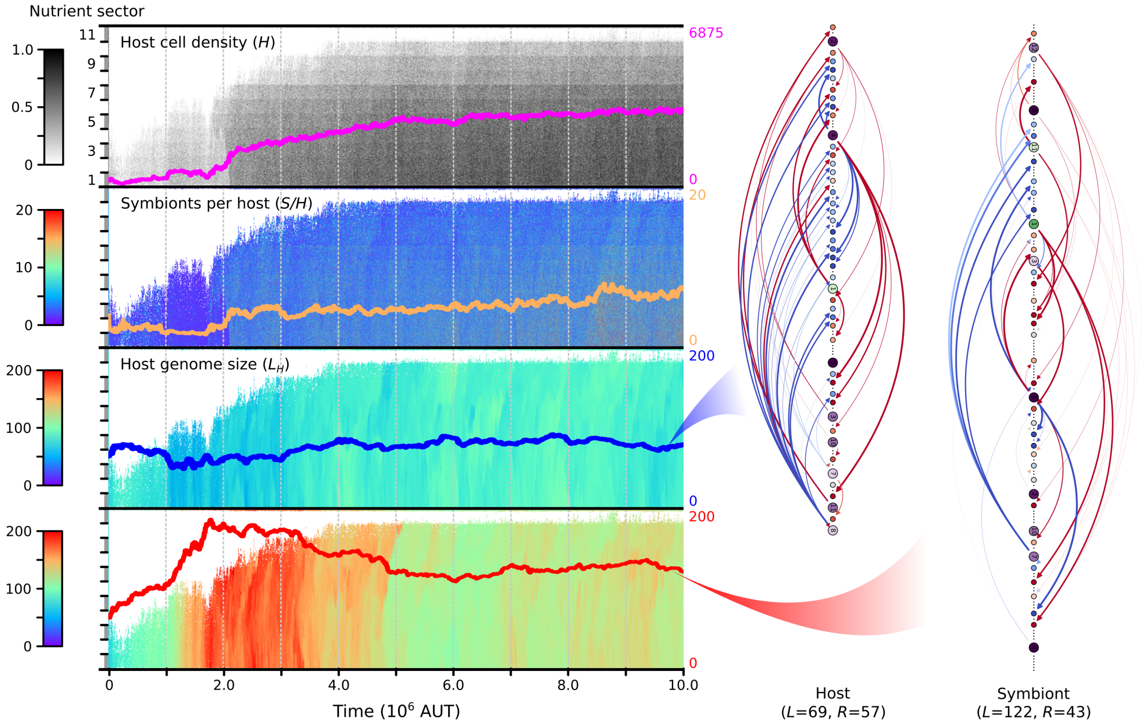


Figure S2: Evolutionary trajectory of P8, i.e. an alternative outcome of evolution from primitive FECA (cf. Fig. 3). Here, holobionts evolve an r-strategy (moderate symbiont numbers) and large symbiont and small host genomes. Symbiont genomes expand dramatically early on when holobionts are still barely viable. The first major adaptation happens at $t \approx 2 \cdot 10^6$, coinciding with an increase in symbiont numbers. Afterwards, symbiont numbers and population size increase only slightly and never reach the levels that characterize the K-strategy as seen in P9. Interestingly, the regulatory repertoire of the host remains larger than that of the symbiont, suggesting that more complex regulation is generally required for the host than for the symbiont, even when a small host genome size is favored.

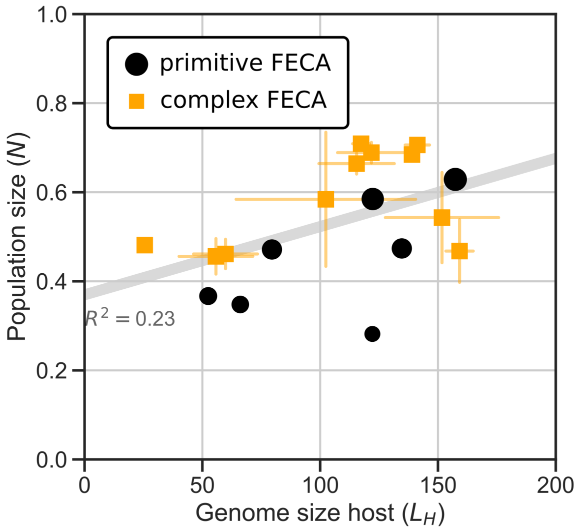


Figure S3: Holobionts with larger host genomes, i.e. eukaryote-like holobionts, have adapted more successfully and achieved greater population size ($p = 0.046$, $N = 18$). For the experiment initialised with complex FECA, 2-3 technical replicates have been averaged and error bars show standard deviations in both dimensions. For the experiment initialised with primitive FECA, the size of the markers is scaled by final population size.

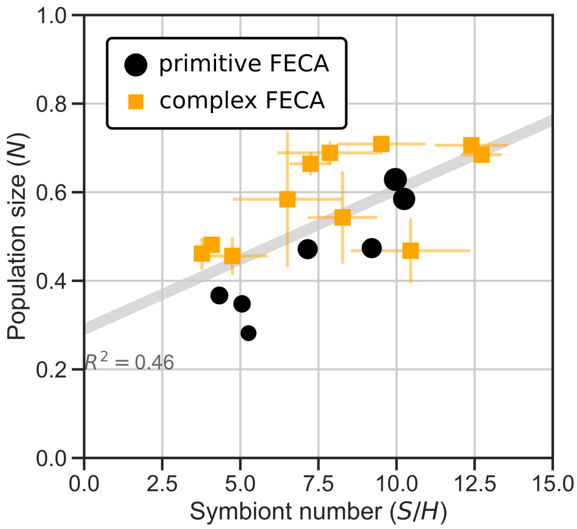


Figure S4: Final population size ($t = 10^7$) correlates with symbiont number across evolution replicates all replicates ($p = 0.0019$, $N = 18$). For the experiment initialised with complex FECA, 2-3 technical replicates have been averaged and error bars show standard deviations in both dimensions. For the experiment initialised with primitive FECA, the size of the markers is scaled by final population size.

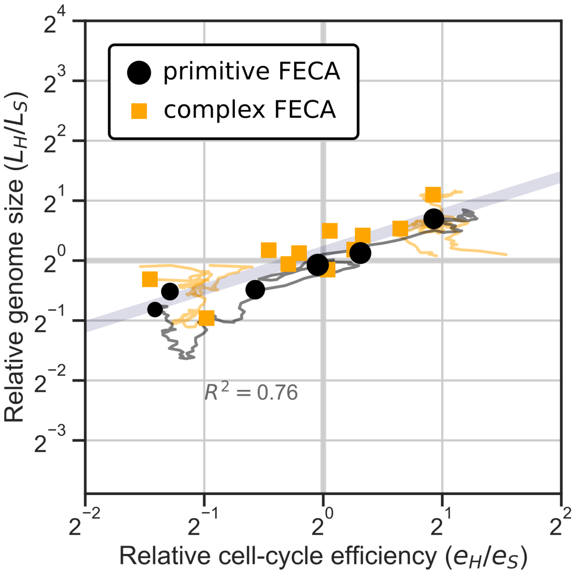


Figure S5: Asymmetry in cell-cycle efficiency yields genome size asymmetry, even when metabolic genes cannot be shared between host and symbiont ($p = 4.43 \cdot 10^{-6}$, $N = 17$). In the presence of non-adaptive forces alone, genome size asymmetry is much less extreme (cf. Fig. 6). The markers show the final timepoint of the experiment. Here, only one technical replicate is shown for each of the replicates initialised with complex FECA. For the experiment initialised with primitive FECA, the size of the markers is scaled by final population size.
